# The effects of terminal tagging on homomeric interactions of the sigma 1 receptor

**DOI:** 10.1101/735803

**Authors:** Hideaki Yano, Leanne Liu, Sett Naing, Lei Shi

**Affiliations:** Computational Chemistry and Molecular Biophysics Unit, National Institute on Drug Abuse - Intramural Research Program, National Institutes of Health, Baltimore, Maryland, United States

## Abstract

The sigma 1 receptor (σ1R) has been implicated in cancers, neurological disorders, and substance use disorders. Yet, its molecular and cellular functions have not been well-understood. Recent crystal structures of σ1R reveal a single N-terminal transmembrane segment and C-terminal ligand-binding domain, and a trimeric organization. Nevertheless, outstanding issues surrounding the functional or pharmacological relevance of σ1R oligomerization remain, such as the minimal protomeric unit and the differentially altered oligomerization states by different classes of ligands. Western blot (WB) assays have been widely used to investigate protein oligomerizations. However, the unique topology of σ1R renders several intertwined challenges in WB. Here we describe a WB protocol without temperature denaturization to study the ligand binding effects on the oligomerization state of σ1R. Using this approach, we observed unexpected ladder-like incremental migration pattern of σ1R, demonstrating preserved homomeric interactions in the detergent environment. We compared the migration patterns of intact σ1R construct and the C-terminally tagged σ1R constructs, and found similar trends in response to drug treatments. In contrast, N-terminally tagged σ1R constructs show opposite trends to that of the intact construct, suggesting distorted elicitation of the ligand binding effects on oligomerization. Together, our findings indicate that the N-terminus plays an important role in eliciting the impacts of bound ligands, whereas the C-terminus is amenable for modifications for biochemical studies.

## 1. Introduction

The sigma 1 receptor (σ1R) is a structurally unique transmembrane protein found in the endoplasmic reticulum (ER) and is associated with intracellular membranes in which it has been posited to act as a molecular chaperone [1]. Although it is broadly distributed in many different tissues and cell types, it is particularly enriched in the central nervous system [2]. σ1R has been implicated in cancer, neurodegenerative diseases, and psychostimulant abuse, and is thus considered a promising therapeutic target [3]. Despite its physiological and pharmacological significance, the structure-function relationships of σ1R are poorly understood. The recent crystal structures of σ1R in complexes with a variety of ligands provide the foundation for mechanistic elucidation of its function at the molecular level [4; 5]. They shed light on the conformation and homomeric status of σ1R in the membrane environment. However, functional categorizations of σ1R ligands remain challenging [6]. Both the differences in intra- or extracellular environments in various σ1R assays and the lack of functional readouts detecting changes in σ1R itself may have contributed to the difficulty in reaching consensus conclusions. Thus, it begs the need for an assay that directly monitors changes occurring in σ1R.

By focusing on σ1R itself, we have developed a bioluminescence resonance energy transfer (BRET)-based assay to categorize σ1R ligands based on presumed changes in σ1R-σ1R interactions [7]. However, the underlying molecular mechanism could not be directly derived from the BRET signal changes. Instead, the specific oligomerization states were characterized by western blot (WB) [7]. This biochemical approach has the advantages of being capable of quantifying the sizes and intensities of migration bands to provide stoichiometric information on homomeric oligomerization, and of allowing comparison across tightly controlled experimental conditions or different pharmacological treatments. In both BRET and WB, σ1R has been subjected to end tagging. However, studies have shown that a bulky protein tag on the N-terminus yields aberrant localization and function, while the C-terminal protein tagging appeared to be permissive and largely retained its function [8].

In the current study, we investigated the effects of N-and C-terminal tagging on the cellular function of σ1R, in particular its homomeric interactions, in order to evaluate the validity of using such constructs in *in vitro* assays. We optimized a WB protocol to detect ligand-induced σ1R oligomerization and compare the results for terminally tagged and unmodified wildtype (WT) σ1R constructs.

## 2. Materials and methods

### DNA constructs, transfection, and cell culture

HEK293T Δσ1R cells were generated using the CRISPR-Cas9 gene deletion method (Santa Cruz). Human σ1R is tagged in pcDNA3.1 plasmid with Myc, NanoLuciferase (Nluc), or mVenus, either N-terminally or C-terminally in frame without any linker (Myc-σ1R, Nluc-σ1R, σ1R-Myc, σ1R-Nluc, or σ1R-mVenus). All constructs were confirmed by sequence analysis. For western blot and radioligand binding, 5 μg (otherwise noted) of terminally tagged and unmodified σ1R plasmid was transfected using lipofectamine 2000 (Invitrogen) for HEK 293T Δσ1R cells in a 10 cm plate. For drug induced BRET, a constant amount of total plasmid cDNA (15 μg) in 1:24 (donor:acceptor ratio for σ1R-Nluc and σ1R-mVenus) was transfected in HEK 293T Δσ1R cells using PEI in a 10 cm plate. Cells were maintained in culture with Dulbecco’s modified Eagle’s medium supplemented with 10% fetal bovine serum and kept in an incubator at 37°C and 5% CO2. Experiments were performed approximately 48 h post-transfection.

### Western blot

HEK293T Δσ1R cells were grown as reported [7] and transiently transfected with the unmodified σ1R, N-terminally tagged Myc-σ1R, N-terminally tagged Nluc-σ1R, C-terminally tagged σ1R-Myc, or C-terminally tagged σ1R-Nluc in 10 cm plates. After 48 hr of growth, confluent cells were harvested in Hank’s Balanced Salt Solution (HBSS), centrifuged at 900 x g for 8 min, and resuspended in HBSS. The cells were then incubated in 1 μM haloperidol, 1 μM PD 144418, 10 μM (+)-pentazocine, or 1% DMSO for 1 h at room temperature. The samples were then centrifuged at 900 x g for 4 min and resuspended in lysis buffer (150 mM NaCl, 1.0% triton X-100, 0.5% sodium deoxycholate, Tris 50 mM, pH 7.5, and protease inhibitors (Roche, catalog# 11697498001)). The samples were sonicated, incubated on ice for 30 min, and centrifuged at 20,000 x g for 30 min. Supernatants were transferred to new tubes. Protein concentrations of the supernatants were determined with Bradford protein assay (Bio-rad, Hercules, CA). Supernatants were mixed with 4x β-mercaptoethanol Laemmli sample buffer to a final 25 μg protein/sample. Samples were electrophoresed on 10% polyacrylamide Tris-glycine gels (Invitrogen) with running buffer (25 mM Tris, 192 mM glycine and 0.1% SDS, pH 8.3, Invitrogen). Proteins were transferred to PVDF membranes (Invitrogen, catalog# IB24002) and immunoblotted with antibodies. anti-GAPDH or anti-actin was used as a loading control. The product information and dilutions of primary and secondary antibodies used are summarized in Supplementary Table 1. Blots were imaged using Odyssey LI-COR scanner and analyzed with LI-COR Image Studio™.

### Radioligand binding assay

Membrane fraction of HEK293T Δσ1R cells was prepared as previously described [7]. The radioligand incubation was carried out in 96-well plates containing 60 μL fresh Earle’s Balanced Salts Solution (EBSS) binding buffer (8.7 g/l Earle’s Balanced Salts without phenol red (US Biological) and 2.2 g/L sodium bicarbonate, pH to 7.4), 20 μL of (+)-pentazocine (varying concentrations), 100 μL membranes (25 μg/well total protein), and 20 μL of radioligand diluted in binding buffer (1 nM [^3^H]-(+)-pentazocine (American Radiolabeled Chemicals), final concentration) for each well. Concentrations for non-radioactive (+)-pentazocine were: 1 mM, 100 μM, 10 μM, 1 μM, 100 nM, 10 nM, 1 nM, and 0.1 nM in EBSS with 10% DMSO. Total binding was determined with 1% DMSO vehicle (final concentration). All compound dilutions were tested in triplicate, and samples were incubated for 120 min at room temperature. The reactions were terminated by filtration through Perkin Elmer Uni-Filter-96 GF/B, presoaked in 0.05% PEI for 120 min, and the 96-well filter plates were counted in Perkin Elmer MicroBeta Microplate Counter as described [7] with counter efficiency at 31% for [^3^H]-(+)-pentazocine. K_d_ and B_max_ values were determined from at least three independent experiments.

### Bioluminescence resonance energy transfer (BRET) assay

Drug-induced BRET is conducted as reported previously [9]. Briefly, cells were prepared in 96-well plates as in acceptor-saturating BRET. 5 μM coelenterazine h was added to each well. Three minutes after addition of coelenterazine h (Nanolight), ligands [(+)-pentazocine (Sigma), PD 144418 (Tocris), and haloperidol (Tocris)] were added to each well in serial dilution. BRET was measured as a ratio between measurements at 535 nm for fluorescence and at 485 nm for luminescence using a Pherastar FSX reader (BMG). Results are calculated for the BRET change (BRET ratio for the corresponding drug minus BRET ratio in the absence of the drug). E_max_ values are expressed as the basal subtracted BRET change in the dose-response graphs.

## 3. Results

### The unmodified WT shows distinct bands up to densities corresponding to octamers

In this study, we used an antibody-based western blot (WB) approach to detect the configurational changes of σ1R oligomerization in response to ligand binding and to the modification of the N- or C-terminus. To exclude the confounding effects of endogenous σ1R, we heterologously expressed (i.e., rescued) various σ1R constructs in σ1R-null (Δσ1R) HEK293T cells (see Methods).

In WB assays, samples would typically be heated, often boiled, in a reducing buffer condition to allow efficient epitope interaction with an antibody in the following step. Indeed, the samples in previous WB assays of σ1R have also been subjected to heating [10; 11; 12], unless using the native gel approach [7]. In this study, we found that by keeping samples on ice before running them on sodium dodecyl sulfate–polyacrylamide gel electrophoresis (SDS-PAGE), we could observe homomeric σ1R interactions when using >5 μg plasmid DNA per 10-cm plate in transfection (Supplementary Figure 1D). Compared to samples receiving boiling treatment (100 °C) that resulted in a single monomer band (∼25 kDa), the samples kept on ice showed a unique ladder-like pattern that corresponds to distinct bands from a monomer up to an octamer (∼200 kDa) of σ1R in WB (Supplementary Figure 2A). As expected, none of the bands were observed in the untransfected Δσ1R cells (Figure 1A). Therefore, we used the on-ice condition (4 °C) for the rest of this study unless otherwise noted.

**Figure 1.**
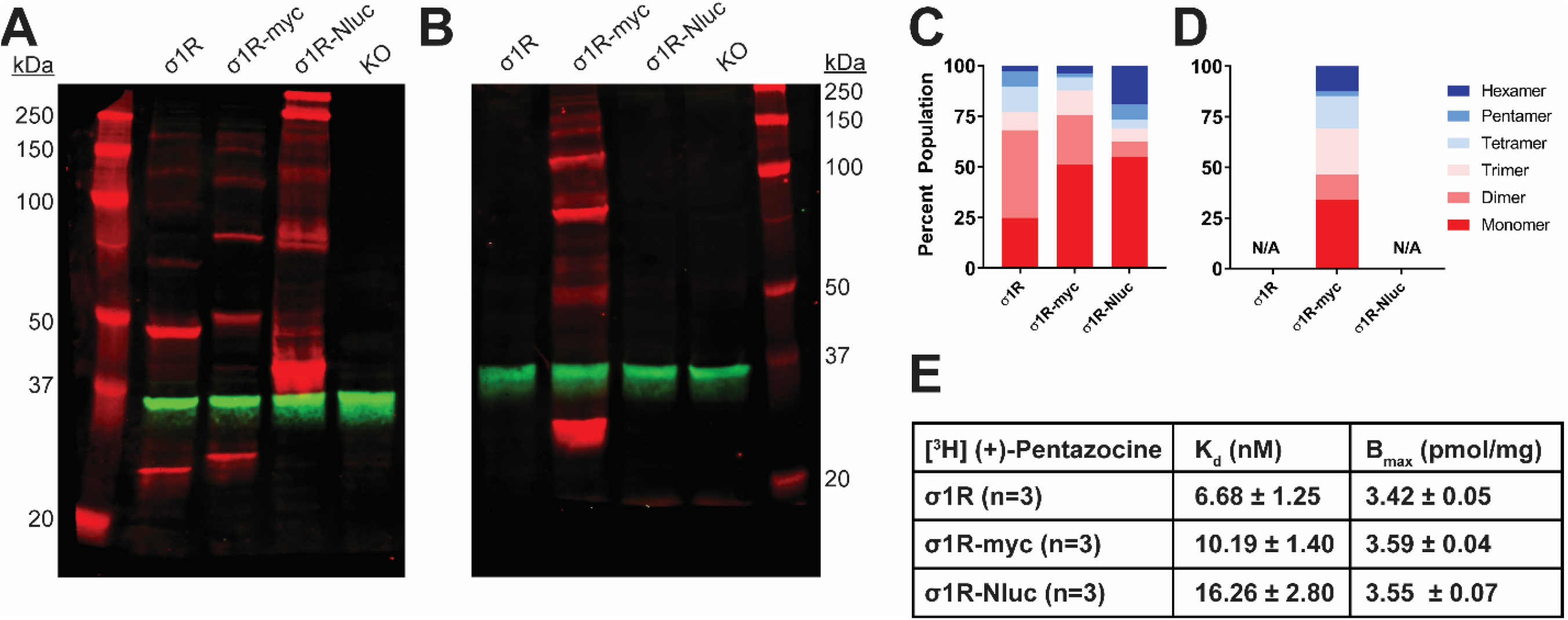
SDS-PAGE mobility comparison among the unmodified and C-terminally tagged σ1R constructs. **A**-**B**. Red bands are visualized by anti-σ1R (**A**) or anti-myc (**B**) antibodies on 10% SDS-PAGE (protein standard, σ1R, σ1R-myc, σ1R-Nluc, σ1R KO cells). Anti-GAPDH bands are shown in green for equal loading. **C-D**. Quantification of band intensities normalized to total signals within each lane for σ1R (**C**) and myc (**D**) blots. The intensities of bands up to hexamer were normalized to the total summation of the densities within the same lane. Due to a close migration pattern in a 10% acrylamide gel, higher order bands past hexamer were not resolvable for quantifications. **E**. Radioligand binding properties of [^3^H] (+)-pentazocine for σ1R, σ1R-myc, and σ1R-Nluc.

In addition to the B-5 (Santa Cruz) σ1R antibody, we chose three other commercially available σ1R antibodies based on their disparate target epitopes (Supplementary Figure 1). We compared the capabilities of these antibodies to detect distinct σ1R oligomer bands in our assay conditions. Similar to B-5, D4J2E (Cell Signaling; Supplementary Figure 1C) antibodies also showed a unique ladder-like pattern but not as distinct as B-5.

### The C-terminally tagged constructs show distinct oligomer bands as well

Previous reports suggest that C-terminal tagging of σ1R does not interfere with localization and function of σ1R [8; 12; 13]. In addition to our previous σ1R construct used in the BRET assay in which Nluc is attached to the C-terminus (σ1R-Nluc) [7], to test the C-terminal tagging’s effect on the σ1R-σ1R interaction, we also generated the σ1R-myc construct, which has a shorter tag at the C-terminus. Both σ1R-Nluc and σ1R-myc exhibit ladder-like patterns similar to the unmodified WT σ1R; the bands range from ∼26 kDa (monomer) to ∼208 kDa (octamer) for σ1R-myc and ∼44 kDa (monomer) to ∼352 kDa (octamer) for σ1R-Nluc (Figure 1A). To confirm the same mobility and density of the bands visualized by an antibody targeting a different epitope, the anti-myc antibody was used against σ1R-myc. In this WB, the bands are generally brighter, which may indicate a stronger epitope affinity of the anti-myc antibody than that of the anti-σ1R antibody (Figure 1B).

To quantify the distribution of σ1R in each oligomeric population for each condition, we measured the intensity of each band (see Methods). Similar patterns were observed across the unmodified and C-terminally tagged constructs (Figure 1C-D). Interestingly, the higher-order bands (i.e., tetramer, pentamer, and hexamer), which are larger than the trimer revealed by the σ1R crystal structures [5], are more intense for σ1R-Nluc and σ1R-myc than the unmodified construct (Figure 1C).

To confirm that the integrity of the ligand binding site and likely the C-terminal domain of the σ1R constructs are not affected by tagging or the WB condition, radioligand binding assay was conducted with [^3^H](+)-pentazocine. Under the same conditions as the WB (i.e., transfection and protein input), the unmodified and C terminally-tagged constructs have virtually the same expression levels and only subtly different K_d_ (Figure 1E).

### The unmodified WT and C-terminally tagged constructs show similar extents of pharmacological changes in higher-order bands

Previously, we have found that σ1R undergoes ligand-induced changes of the oligomerization states [7]. Here, we investigated the effect of ligand binding on the migration pattern in WB. σ1R ligands were added to σ1R-expressing cells and incubated for 1h before proceeding to sample preparation. For unmodified WT σ1R, both haloperidol and PD144418 decreased the intensities of the higher-order bands (i.e., tetramer, pentamer, and hexamer), while (+)-pentazocine increased their intensities (Figure 2A-C). These results are consistent with our previous findings using BRET assays that categorized haloperidol, PD144418, and 4-PPBP into haloperidol-like ligands and (+)-pentazocine and PRE-084 into pentazocine-like ligands [7]. Consistently, both σ1R-myc and σ1R-Nluc showed similar extents of changes in the higher-order bands, in which haloperidol and PD144418 decreased and (+)-pentazocine increased the relative densities of the higher-order bands (Figure 2D-I). The opposing effects of these σ1R ligands are consistent with our previous study [7].

**Figure 2.**
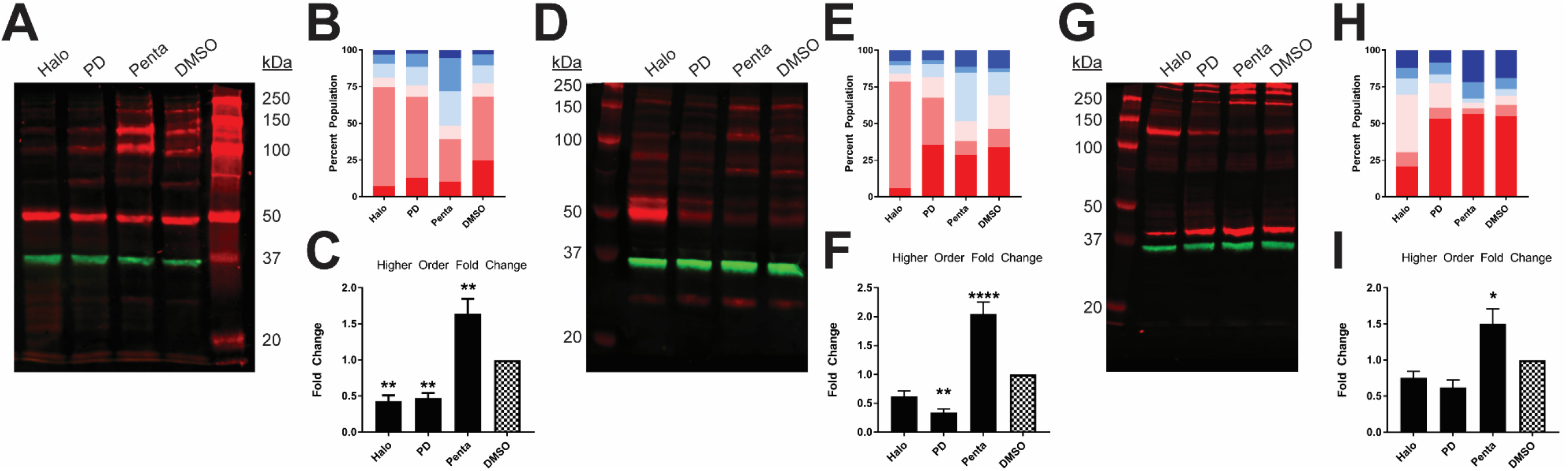
Pharmacological effects on SDS-PAGE mobility among the unmodified and C-terminally tagged σ1R constructs. Results for the unmodified (**A**-**C**), C-terminally myc-tagged (**D**-**F**), and C-terminally Nluc tagged (**G**-**I**) constructs are shown. **A**, **D**, **G**. Red bands are visualized by anti-σ1R (**A**, **G**) or anti-myc (**D**) antibodies and green bands are visualized by anti-GAPDH (**A**, **D**, **G**) antibodies on 10% SDS-PAGE (protein standard, haloperidol, PD 144418, (+)-pentazocine, DMSO vehicle). **B**, **E**, **H**. Quantification of band intensities normalized to total signals within each lane for σ1R (**B**, **H**) and myc (**E**) blots. **C**, **F**, **I**. Pharmacological changes in higher order densities corresponding to tetramers, pentamers, and hexamers normalized to DMSO vehicle (haloperidol, PD 144418, (+)-pentazocine, DMSO vehicle).

In contrast to samples prepared on ice, temperature increases in the sample buffer decreased the densities of the dimer and above bands, indicating that σ1R-σ1R interaction is disrupted by the temperature change (Supplementary Figure 2A). In the presence of different ligands, the 70°C treatment completely monomerized σ1R as well, while the monomer densities across different ligand treatments are similar (Supplementary Figure 2B). Thus, the σ1R expression level is unlikely changed by the ligands, because changes in the total copy number would affect monomer band density.

### The N-terminally tagged myc-σ1R and Nluc-σ1R exhibit very weak higher order bands

As opposed to the C-terminal tagging, it has been reported that N-terminal tagging of σ1R interferes with the localization of σ1R [8]. To study the effect of N-terminal tagging on gel mobility, in addition to our previously studied Nluc-σ1R [7], we generated the myc-σ1R construct. Compared to the unmodified WT construct, myc-σ1R showed significantly fainter bands for trimer and higher-order bands. In both myc-σ1R and Nluc-σ1R, the higher-order bands have much less relative distributions than the unmodified construct (Figure 3A,C). When incubated with anti-myc antibody, the fraction of total density that was present in higher-order bands for myc-σ1R did not notably improve (Figure 3B, D). Nluc-σ1R also showed reduced intensities in higher-order bands, which is more apparent when compared to the density distribution of the C-terminally tagged σ1R-Nluc (Figure 3A,C and Figure 1A,C). Even though the oligomerization states are drastically different from their corresponding C-terminal tagged constructs, radioligand binding results showed K_d_ and B_max_ are not significantly different from those of the unmodified construct (Figure 3E). Therefore, the significantly decreased relative densities of higher-order bands in WB are likely due to changes in receptor conformation and epitope exposure by the N-terminal tagging.

**Figure 3.**
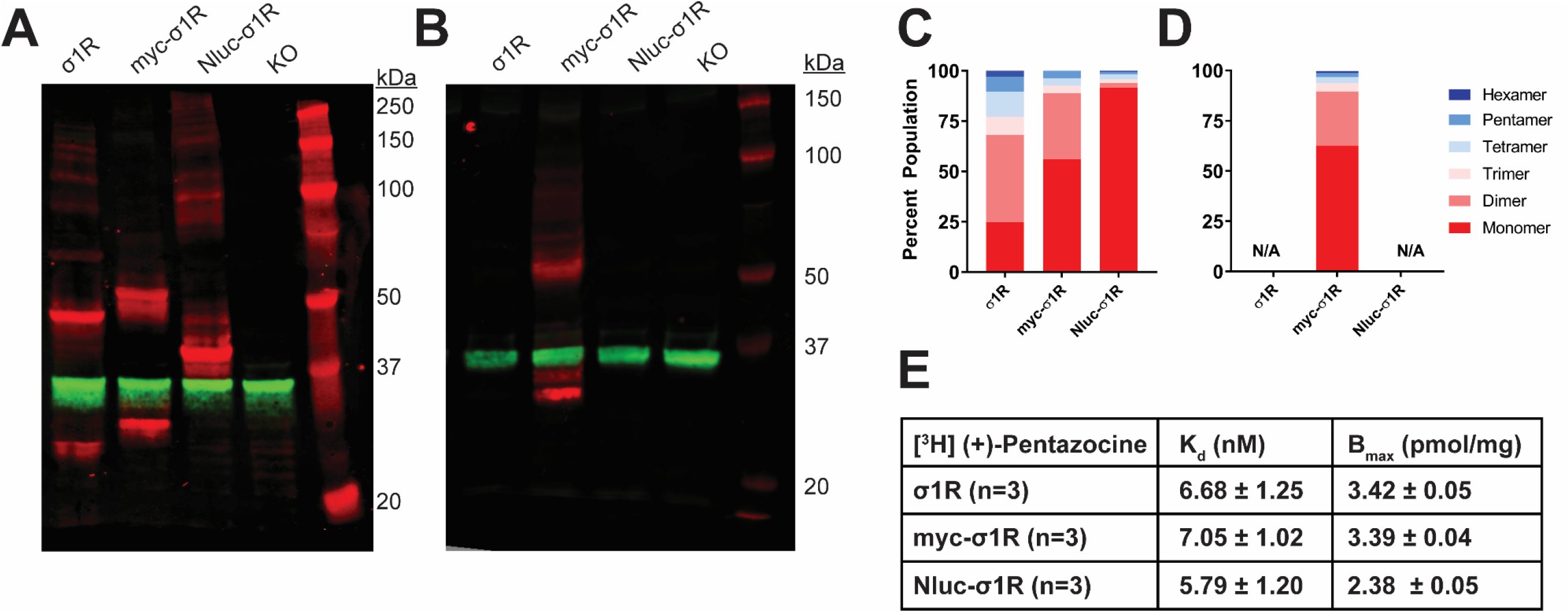
SDS-PAGE mobility comparison among the unmodified and N-terminally tagged σ1R constructs. **A-B**. Red bands are visualized by anti-σ1R (**A**) or anti-myc (**B**) antibodies and green bands are visualized by anti-GAPDH (**A**-**B**) antibodies on 10% SDS-PAGE (protein standard, σ1R, myc-σ1R, Nluc-σ1R, σ1R KO cells). **C-D**. Quantification of band intensities normalized to total signals within each lane for σ1R (**C**) and myc (**D**) blots. **E**. Radioligand binding properties of [^3^H] (+)-pentazocine for σ1R, myc-σ1R, and Nluc-σ1R.

### The N-terminally tagged constructs show substantially different pharmacology from unmodified WT σ1R

Next, we studied the pharmacology of myc-σ1R and Nluc-σ1R in gel mobility to assess the effects by short and long N-terminal tagging. For myc-σ1R, haloperidol increased the higher-order bands while it diminished the monomer band (Figure 4A-C). PD144418 did not affect the relative density of each band, compared to the DMSO treated sample. (+)-pentazocine decreased the intensities of higher-order bands while increasing that of the monomer band. For Nluc-σ1R, haloperidol increased the higher order bands slightly while PD144418 showed effectively no change. (+)-pentazocine showed an increased monomer band (Figure 4D-F). Overall, there are similarities between myc-σ1R and Nluc-σ1R for their pharmacology. Haloperidol increased the densities of higher-order bands, while (+)-pentazocine increased that of monomer bands. However, these pharmacological trends are opposite to that of the unmodified σ1R (Figure 2A-C), indicating that the N-terminal tagging may interfere with how ligand binding propagates the conformational changes within σ1R or directly interferes with the formation of oligomers.

**Figure 4.**
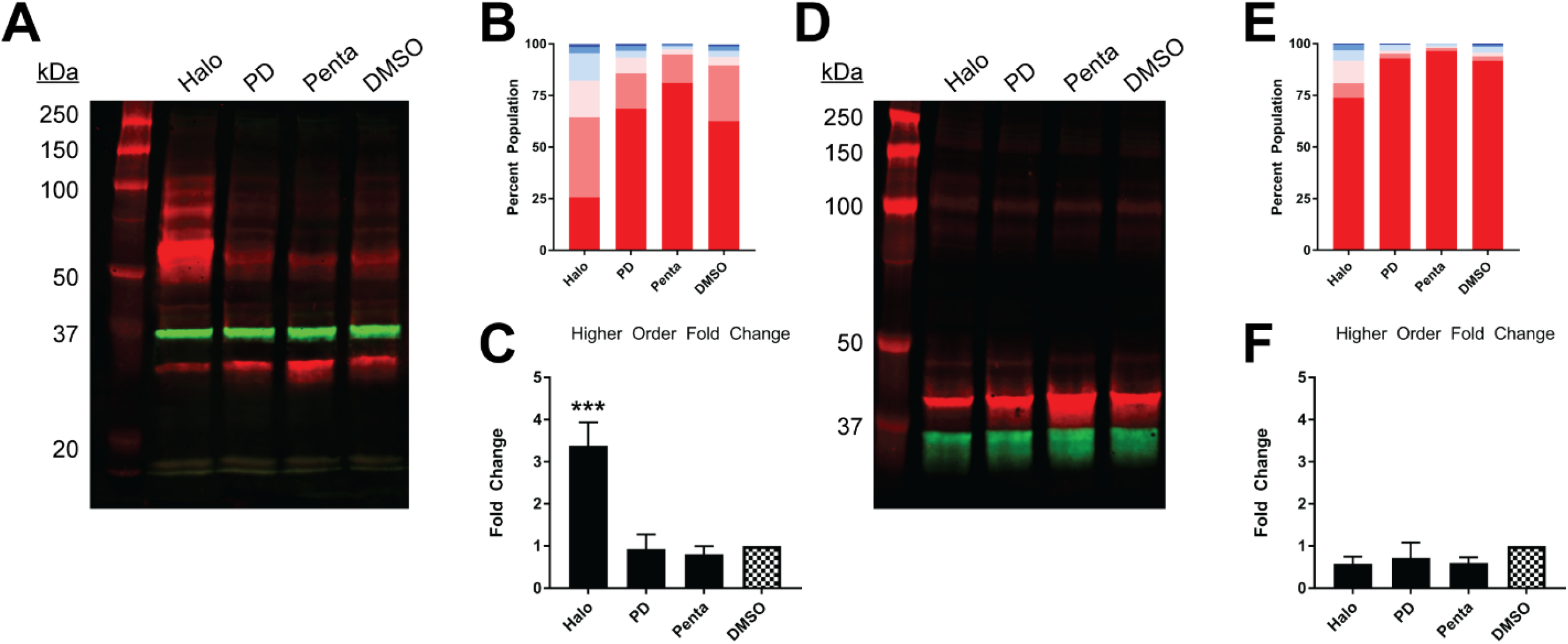
Pharmacological effects on SDS-PAGE mobility of the N-terminally tagged σ1R constructs. Results for the N-terminally myc-tagged (**A**-**C**), and N-terminally Nluc-tagged (**D**-**F**) constructs are shown. **A**, **D**. Red bands are visualized by anti-myc (**A**) or anti-σ1R (**D**) and green bands are visualized by anti-GAPDH (**A**, **D**) antibodies on 10% SDS-PAGE (protein standard, haloperidol, PD 144418, (+)-pentazocine, DMSO vehicle). **B**, **E**. Quantification of band intensities normalized to total signals within each lane for myc (**B**) and σ1R (**E**) blots. **C**, **F**. Pharmacological changes in higher order densities corresponding to tetramers and above bands normalized to DMSO vehicle (haloperidol, PD 144418, (+)-pentazocine, DMSO vehicle).

## 4. Discussion

σ1R has been extensively studied at molecular level using biochemical and pharmacological approaches [10; 13; 14; 15; 16; 17; 18]. In this study we investigated the effects of terminal tagging on σ1R oligomerization and pharmacology. In particular, in conjunction with WB, the end-tagged constructs revealed the ability of N-terminal modification to interfere with the oligomerization of σ1R. SDS-PAGE WB is one of the most commonly used biochemical assays to quantify protein density. Because of detergent solubilization, protein-protein interactions in their native cellular environment can be disrupted in many instances. Here, however, by maintaining samples on ice, we were able to preserve homomeric σ1R-σ1R interactions and visualize discrete bands corresponding to oligomers as large as octamers by using an antibody (B-5) targeting an epitope at residues 136-169 of σ1R. As a technical note, after the antibody has been stored for a long time, the antibody staining becomes dimmer and the higher-order bands cannot be visualized, potentially due to degradation (Supplementary Figure 1E-F). Interestingly, the other antibodies against distinct epitopes from different vendors resulted in different visualization patterns indicating the corresponding epitopes may be differentially exposed in different oligomerization states, while residue 136-169 might always be exposed. Although intriguing, these antibodies were not further pursued because of their significantly lower band intensities. In general, the WB method described herein is deemed useful to study σ1R pharmacology since ligand-induced conformational changes, particularly in higher-order bands, can be quantitatively evaluated.

We have previously reported a BRET-based proximity assay to study σ1R pharmacology in HEK293T cells [7]. The C-terminally tagged σ1R-Nluc construct used in that study showed a similar response to ligands in the migration pattern of higher-order bands as the unmodified σ1R, demonstrating that BRET responses very likely reflect the configurational changes in σ1R oligomerization observed in the unmodified σ1R. In current study, we used Δσ1R HEK293T cells. By repeating the BRET experiment in Δσ1R HEK293T cells transfected with σ1R-Nluc and σ1R-Venus (Supplementary Figure 3), we found the σ1R pharmacology effectively remained the same as the previously reported BRET results in HEK293T cells [7]. Given that in the BRET assays, haloperidol and PD144418 increased BRET signals while in the WB assays, they decreased higher-order band densities and increased lower-order band densities, we hypothesize that the BRET readouts recapitulate the changes in the lower-order populations. In that regard, the dimer and trimer populations comprise 48.9% in haloperidol and 24.3% in PD144418, compared to 13.7% in DMSO (Figure 2H). The pattern can be correlated with the BRET_max_ order, in which haloperidol shows higher BRET_max_ than PD144418 (Supplementary Figure 3).

We observed that the total signal (*i.e.*, summation of band densities up to hexamer) increased with (+)-pentazocine (38%) and decreased with haloperidol (−25%) or PD144418 (−23%) compared to DMSO lane (Figure 2A). Since there is no change in σ1R expression level across different ligand conditions (Supplementary Figure 2B), these changes in total signal, primarily due to changes in the higher-order bands, are caused by changes in either the copy number and/or changes in epitope accessibility in higher-order oligomers. If there is an increase in copy number of higher order bands, the mobilized populations should equal the collective decrease in lower-order bands. However, as we only observed a weak complementary decrease in lower-order bands, we postulate that changes in the epitope affinity in higher-order populations of σ1R plays a role in increased higher-order band intensities.

In addition, we found that the N-terminal modification limits higher-order interactions of σ1R in our solubilization condition, consistent with the previously reported detrimental effects of N-terminal tagging or point mutation on the cellular localization and function [8]. It is noteworthy that there is a putative Arg-Arg ER retention signal [19] at the N-terminus of σ1R. We propose that the steric hindrance of the N-terminally tagged protein may interfere with the recognition of Arg-Arg tandem residues at the 7^th^ and 8^th^ positions. This interference is potentially akin to some other membrane proteins that escape from ER retention due to its Arg-Arg motif masked by nearby domains (e.g., GABA_B_ receptor [20]). Despite its probable mislocalization, we presume that the folding of σ1R should only be modestly affected by the N-terminal tagging because of the preserved binding pocket considering the only lightly-affected K_d_ compared to that of the unmodified construct. Therefore, the reduced σ1R-σ1R interaction may arise from the membrane environment in which σ1R is localized such that the σ1R density may be too low to find an interacting partner. Further, while the exterior of the barrel motif of the C-terminal domain has been revealed by the crystal structures to be the interface for trimer, it is likely other part of the of the protein involves in forming other oligomer states, such as the N-terminus and N-terminal transmembrane helix [21]. The N-terminal tagging may potentially bias the protein towards dimer formation by facilitating the interactions between two monomers at their N-terminal region.

## Author contribution

H.Y. and L.S. designed the study. L.L., S.N., and H.Y. performed the experiments. All authors took part in interpreting the results. H.Y. wrote the initial draft, with L.L. and L.S. participating in revising the manuscript.

## Acknowledgements

We would like to thank Drs. Yoki Nakamura and Tsung-Ping Su at NIDA for collaborating with us to develop Δσ1R HEK293T cells used in this assay and sharing their image detection instrument (LI-Cor). We thank Dr. Andrew Fant for insightful discussions. Support for this research was provided by the National Institute on Drug Abuse–Intramural Research Program, Z1A DA000606-03 (L.S.).

---

**Supplementary Figure 1.**
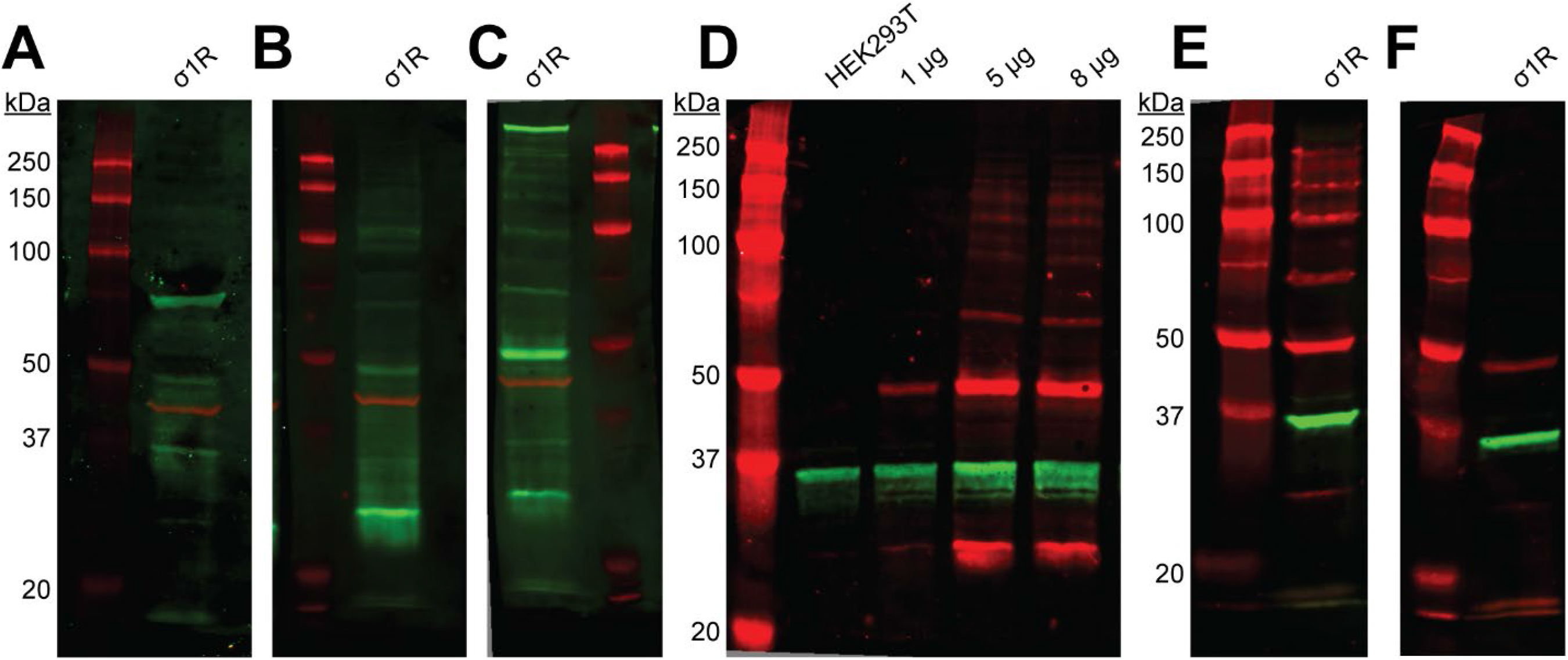
Immuno-reactivities of commercially available σ1R antibodies. **A-C**. Green and red bands are visualized by anti-σ1R (**A**. AB 53852, Abcam, **B**. AB 223702, Abcam, **C**. DHJ2E, Cell Signaling) and anti-actin antibodies respectively on 10% SDS-PAGE. **D**. Endogenous σ1R expressed in WT HEK293T cells or increasing amounts of σ1R expressed in σ1R KO cells (1, 5, 8 μg) are detected by the anti-σ1R antibody (B-5, Santa Cruz) in red along with GAPDH loading control in green. **E-F**. Immuno-reactivities are diminished over a prolonged storage time (**E**. ∼1 month vs. **F**. > 6 months in 4 °C).

**Supplementary Table 1.** Product information and dilutions of primary and secondary antibodies

|  | Product ID, Vendor | Epitope (residue numbers) | Dilution |
| --- | --- | --- | --- |
| *Primary antibody* |  |  |  |
| σ1R (mouse monoclonal) | B-5, Santa Cruz | 136-169 | 1:1,000 |
| σ1R (rabbit polyclonal) | AB 53852, Abcam | C-terminus | 1:625 |
| σ1R (rabbit polyclonal) | AB 223702, Abcam | 35-88 | 1:250 |
| σ1R (rabbit monoclonal) | DHJ2E, Cell Signaling | around 137 | 1:1,000 |
| GAPDH (rabbit monoclonal) | 14C10, Cell Signaling | C-terminus | 1:2,000 |
| Actin (mouse monoclonal) | C-2, Santa Cruz | 357-375 | 1:2,000 |
| Myc (mouse monoclonal) | 9B11, Cell Signaling | 410-419 | 1:1,000 |
| *Secondary antibody* |  |  |  |
| Donkey $\alpha$ mouse | IRDye 680RD, LI-COR | | 1:5,000; 1:10,000 |
| Goat $\alpha$ rabbit | IRDye 800CW, LI-COR | | 1:10,000; 1:5,000 |

**Supplementary Figure 2.**
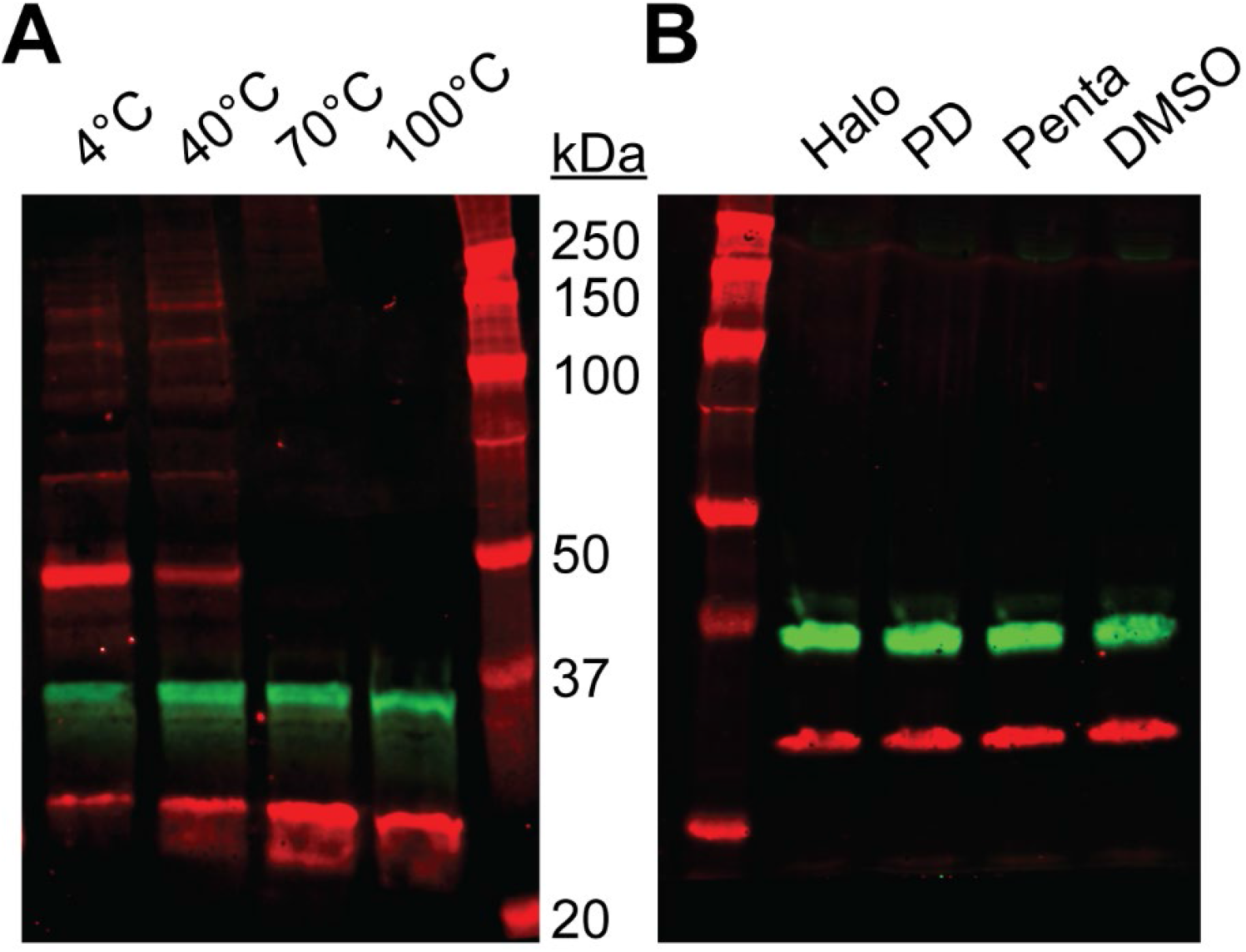
Protein sample preparation at different temperatures. **A**-**B**. Red and green bands are visualized by anti-σ1R and anti-GAPDH antibodies respectively on 10% SDS-PAGE. **A**. Samples were prepared in increasing temperatures (3, 40, 70, and 100 °C). **B**. Ligand-treated samples were heated at 70 °C for 15 min (protein standard, haloperidol, PD 144418, (+)-pentazocine, DMSO vehicle).

**Supplementary Figure 3.**
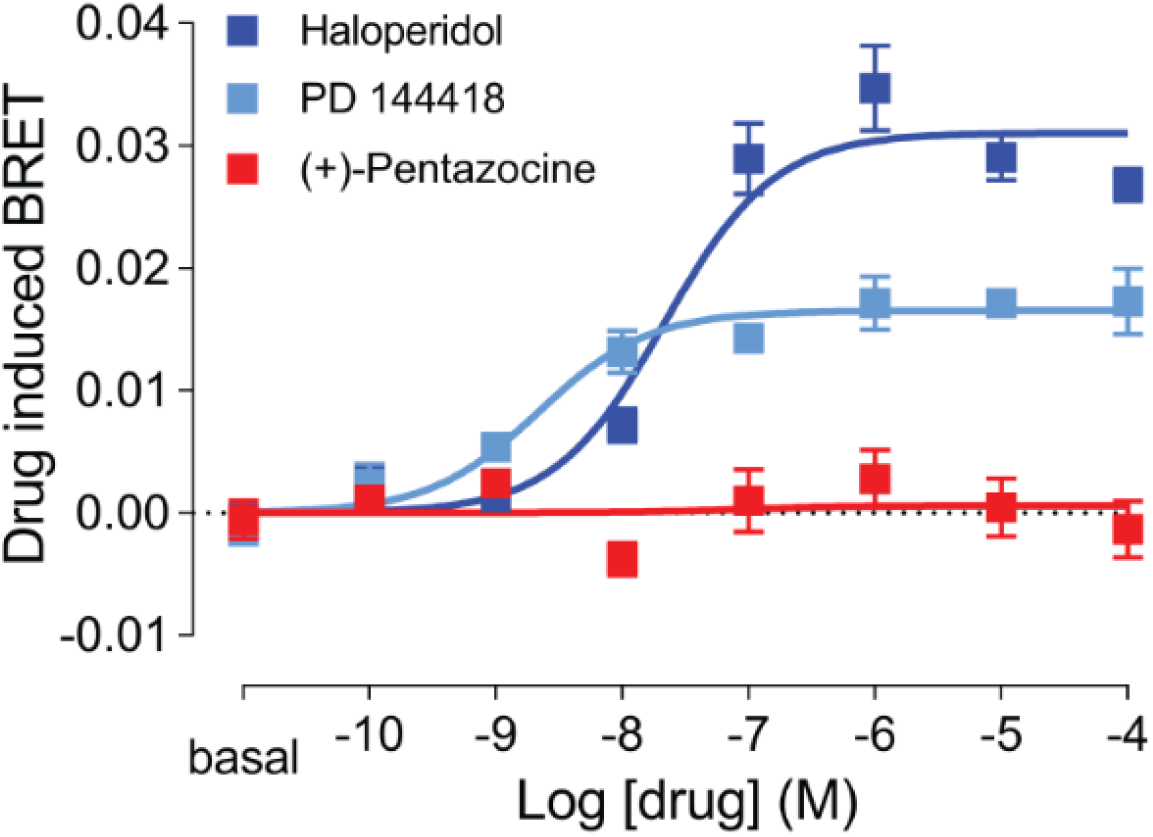
**A**. Drug induced changes of σ1R homomer BRET. Drug induced BRET between the C-terminally tagged σ1R-Nluc and σ1R-Venus in σ1R KO cells is detected at 60 min for (+)-pentazocine (red), haloperidol (blue), and PD144418 (cyan). Data represents mean±S.E.M. (n=5 or more).

